# Facile method for screening clinical T cell receptors for off-target peptide-HLA reactivity

**DOI:** 10.1101/472480

**Authors:** Marvin H. Gee, Xinbo Yang, K. Christopher Garcia

**Affiliations:** Departments of Molecular and Cellular Physiology and Structural Biology, Stanford University School of Medicine, Stanford, CA 94305, USA; Program in Immunology, Stanford University School of Medicine, Stanford, CA 94305, USA; The Howard Hughes Medical Institute, Stanford University School of Medicine, Stanford, CA 94305, USA

## Abstract

T cell receptors (TCRs) exhibit varying degrees of cross-reactivity for peptides presented by the human leukocyte antigen (HLA). In engineered T cell therapies, TCR affinity maturation is a strategy to improve the sensitivity and potency to often a low-density peptide-HLA (pHLA) target. However, the process of affinity maturation towards a known pHLA complex can introduce new and untoward cross-reactivities that are difficult to detect and raises significant safety concerns. We developed a yeast-display platform of pHLA consisting of ~100 million different 9mer peptides presented by HLA-A*01 and used a previously established selection approach to validate the specificity and cross-reactivity of the A3A TCR, an affinity-matured TCR against the MAGE-A3 target (EVDPIGHLY). We were able to identify reactivity against the titin peptide (ESDPIVAQY), to which there is now known clinical toxicity. We propose the use of yeast-display of pHLA libraries to determine cross-reactive profiles of candidate clinical TCRs to ensure safety and pHLA specificity of natural and affinity-matured TCRs.

## INTRODUCTION

Recent advances in T cell therapies have resulted in significant improvements in anti-tumor responses when treating various cancer types (1). These therapies utilize the adoptive transfer of T cells specific for a known cancer antigen, typically a self-antigen known to be overexpressed or expressed exclusively on the tumor (2,3). Although the types of antigens being targeted have significantly shifted towards neoantigens, tumor-specific mutations that can be immunogenically distinct from its wildtype counterpart, the concept of adoptive T cell transfer remains the same (4).

Regardless of the antigens targeted, a major focus in T cell therapies is developing TCRs with high levels of on-target specificity and preventing off-target toxicity due to TCR cross-reactivity. In some cases, TCRs may simultaneously recognize the wild-type antigen and corresponding neoantigen (5,6), which can deter the use of a particular TCR. When targeting wild-type antigens, off-target cross-reactivity has been a significant issue in several clinical trials and has resulted in patient deaths (7–9), despite extensive testing for pre-clinical toxicity.

Intuitively, the benefit of TCR cross-reactivity is to enable host protection against pathogens and to reduce the burden for TCR diversity and T cell number to cover the diversity of targetable peptides (10–12). Despite this, the diversity of the repertoire is still significant with approximately 10^12^ T cells in the human body with a diversity of 10^7-8^ unique receptors (13). Recent studies indicate that TCRs recognize peptides on the order of thousands to tens of thousands, in which the TCR contact residues on the peptide are highly restricted in sequence, while the non-contacting peptide residues can exhibit sequence drift (14). While TCRs generally appear highly specific for their endogenous target, in addition to peptides that bear close sequence resemblance (14,15), cross-reactivity can occur, and be attributed to many different factors including conformational flexibility of the TCR complementarity-determining regions (16), alternative TCR ‘footprints’ contacting the pHLA (17), and molecular mimicry (18).

Although many studies have characterized the features of TCR cross-reactivity (19), there exists limited high-throughput methodologies to comprehensively measure the cross-reactivity of candidate TCRs (20). One such methodology is the use of yeast-display of pHLA libraries to survey the landscape of peptides that can be bound by a TCR of interest (14,15,21). These libraries display on the order of hundreds of millions of peptides covalently linked to a single HLA or MHC allele. A recombinantly-expressed soluble TCR can be used to physically isolate yeast expressing pHLA ligands. After several rounds of selection, the yeast culture can be deep-sequenced to identify the peptide landscape that can be recognized by a TCR and algorithms can be used to determine human and/or other organismal specificities. This process can be repeated for any number of TCRs and is amenable to high-throughput.

In order to determine if pHLA libraries can be used to accurately characterize the crossreactivity of a TCR, we developed the HLA-A*01 yeast-display library to test the clinically-used A3A TCR, previously tested in the clinic to treat myeloma and melanoma (8). The A3A TCR is known to recognize the MAGE-A3 antigen EVDPIGHLY, but was also found to be crossreactive with the Titin antigen ESDPIVAQY resulting in patient deaths, and this off-target specificity was identified only following retrospective analysis of patient samples (7,8).

## RESULTS

### Peptide-HLA-A*01 library composition and TCR selections

We first generated the HLA-A*01 library using the same scaffold as previously used for HLA-A*02:01, in which the peptide is N-terminally linked via a flexible (Gly4-Ser)3 linker to human β2M (Fig. 1A) (14). The β2M is followed by the HLA-A*01 heavy chain and followed by an epitope tag (c-Myc) and the yeast mating protein Aga2. The expression vector utilizes the native Aga2p:Aga1p interaction to display proteins on the surface (22) and the epitope tag used to monitor pHLA expression over rounds of selection. We generated a 9mer peptide library for HLA-A*01 using HLA anchors at P3 and P9 restricted to D/E and Y, respectively, based on known peptides presented by HLA-A*01 (Fig. 1A) (23). We randomized the remaining positions to maximize the available diversity of the library to recover binding peptides for a given TCR. The library construction yielded a functional library size of 180 million total yeast clones determined by limiting dilution and counting of yeast colonies.

**Figure 1.**
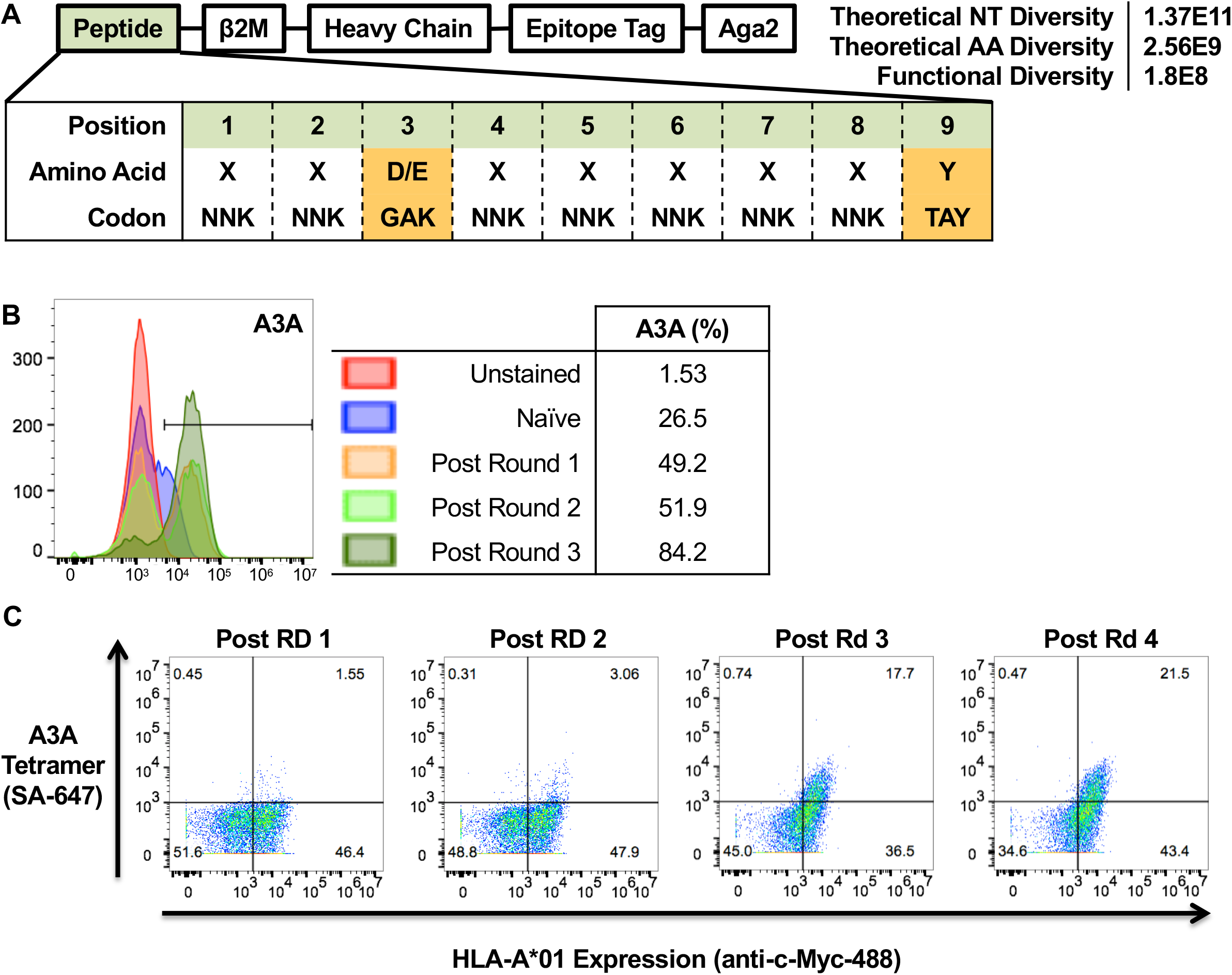
HLA-A*01 library and selection by the A3A TCR. A) Design of the HLA-A*01 library construct for yeast-display from N-to C-terminus of the displayed protein. An epitope tag is introduced to monitor protein induction during the selection process. The theoretical nucleotide and amino acid diversity are calculated from the library composition. Peptide anchors are marked in orange and are limited to one or two amino acids. Codons follow the IUPAC naming system. B) c-Myc induction of the yeast-display library measured by anti-c-Myc-488 antibody through the rounds of selection. C) TCR tetramer at 400 nM and anti-c-Myc-488 co-staining of the yeast after re-growth and induction of all rounds.

We performed yeast selections with 4 rounds of recombinantly expressed A3A TCR identified previously to be specific for MAGE-A3 presented by HLA-A*01 (8). Through the selection process, an antibody specific for the c-Myc epitope tag co-expressed with HLA-A*01 was used to quantify expression of the peptide-HLA-A*01 library, showing increasing amounts of protein display over time (Fig. 1B). The A3A TCR selection showed an increasing trend in total percentages of yeast expressing c-Myc, indicating an enrichment of yeast clones bearing pHLA that may bind the A3A TCR (Fig. 1B). After all rounds of selection were completed, the selected pool of yeast from each round were grown and induced to express pHLA and tetramer stained with the A3A TCR (Fig. 1C). The yeast selected by the A3A TCR showed positive TCR tetramer staining by rounds 3 and 4 of the selection, indicating enrichment of yeast clones that displayed peptides binding to the A3A TCR. This correlated well with the percentage of yeast expressing the c-Myc epitope tag per round (Fig. 1B).

### The A3A TCR selects a converged set of peptides resembling that of MAGE-A3

The pool of yeast was then deep-sequenced for each round of the selection. The yeast clones selected by the A3A TCR converged to a set of 70 relatively related peptides at round 3 (Fig. 2A-C). When accounting for the abundance of the unique peptides by counts from deep-sequencing, the amino acids per position resemble that of the MAGE-A3 peptide (Fig. 2A). When observing the amino acids that are present in all 70 of the unique peptides, without accounting for enrichment by sequencing, there was complete amino acids representation for all positions of the MAGE-A3 peptide and only one missing at P8 for the Titin peptide (Fig. 2B). A word logo plot of the unique peptides at round 3 of the selections reveal semblance to the MAGE-A3 peptide and Titin and highlight ‘hotspots’ of TCR recognition, such as the preference for Pro at P4 (Fig. 2C). Disregarding the relatively fixed anchor residues, the selected peptides resembled that of the MAGE-A3 peptide at several positions including P1, P2, P4, P5, and P7. Interestingly, the peptide landscape reveals a strong preference of limited amino acids of the A3A TCR for the N-terminus as opposed to the C-terminus. Altogether, the selected peptide data reveals a semblance to the MAGE-A3 and Titin peptides, which may be explained by more tolerant recognition at the C-terminal end.

**Figure 2.**
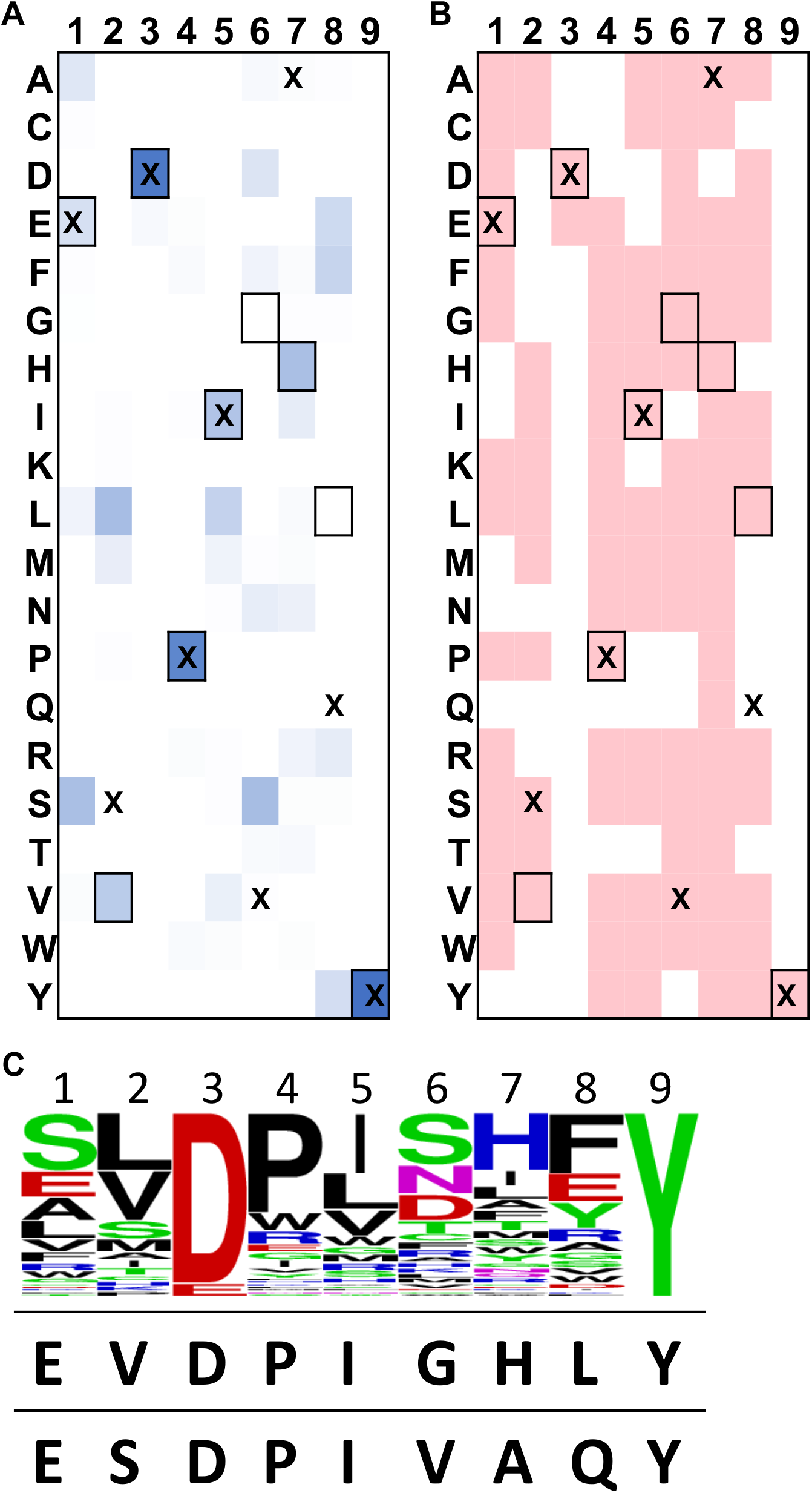
Selection results of the A3A TCR on the HLA-A*01 library. A) Heatmap identifying the amino acid composition of peptides isolated in round 3 of the selection accounting for enrichment by sequencing and B) binary presence of an amino acid from the 70 unique peptides. Outlined boxes are the amino acids of the MAGE-A3 peptide EVDPIGHLY. Boxes that contain an “X” indicate amino acids of the Titin peptide ESDPIVAQY. C) Word logo generated from the unique peptides isolated in round 3 of the selection disregarding the deep-sequencing round counts. The height of the amino acids represents the proportion of abundance of those amino acids among the 70 unique peptides. The MAGE-A3 peptide and titin peptide are listed below the word logo, respectively.

## DISCUSSION

Adoptive T cell transfer has shown promising results in clinical trials with cancer patients (1); however, there are still concerns over off-target toxicities due to TCR cross-reactivities for self-antigens that have arisen as a result of affinity maturation (8,9). Developing a reliable high-throughput methodology to determine the targets and cross-reactivities of TCRs is attractive for monitoring the safety of clinical candidates ahead of clinical studies. pHLA libraries have been previously used to successfully characterize TCR specificities and cross-reactivities (15). The technology described here could provide an avenue to ‘measure’ the cross-reactivity of multiple wild-type or engineered TCRs for a given target pHLA. Such an approach could streamline the identification of candidate TCRs for further clinical development based on antigen specificity profiling as an indirect readout of clinical safety.

Here, we show the practicality of using pHLA libraries to characterize the peptide landscape recognized by each TCR and how this information can be used to identify off-target specificities of lead TCR compounds that may introduce clinical toxicity.

## METHODS

### T cell receptor expression

T cell receptors were expressed as described previously (14).

### Generating the HLA-A*01 library

The HLA-A*01 library was generated similarly to the scaffold used to display the HLA-A*02:01 library system (14). The heavy chain also contained a Y84A mutation to provide an opening for the C-terminus of the peptide to be covalently linked to β2M by an engineered linker. We generated a 9mer peptide library for HLA-A*01 using HLA anchors at P3 and P9 restricted to D/E and Y, respectively (23) (Fig. 1A). The library was generated as described previously using amplification of a parent stop-template vector with degenerate codons (14). A limiting dilution of the initial yeast library at 1:10,000, 1:1,000, 1:100, and 1:10 yielded a functional library size of 1.8x 10^8^ total yeast clones.

### Selecting the HLA-A*01 library

The HLA-A*01 library was selected as done previously utilizing streptavidin-coated magnetic beads (14). Tetramer was generated at 400 nM using a 5:1 mixture of TCR monomer to multivalent streptavidin linked to 647. Yeast were stained at 100,000 cells in 200 μL using a 1:100 anti-c-Myc-488 antibody (9402S, Cell Signaling).

### Deep-sequencing

The HLA-A*01 library was deep sequenced and processed as done previously (14) using the Illumina MiSeq.

## ACKNOWLEDGEMENTS

M.H.G. was supported by F31 CA216926-01. MHG was supported by a Stanford Graduate Research Fellowship and F31 CA216926-01. KCG was supported by AI103867. We would like to acknowledge the Parker Institute for Cancer Immunotherapy and the Howard Hughes Medical Institute for supporting the research. MHG and KCG are co-founders in 3T Biosciences, Inc.

## AUTHOR CONTRIBUTIONS

MHG and XY contributed to experimental design and execution. KCG supervised all aspects of the study.

